# Meta-analysis of liver and heart transcriptomic data for functional annotation transfer in mammalian orthologs

**DOI:** 10.1101/123414

**Authors:** Pía Francesca Loren Reyes, Tom Michoel, Anagha Joshi, Guillaume Devailly

## Abstract

Functional annotation transfer across multi-gene family orthologs can lead to functional misannotations. We hypothesised that co-expression network will help predict functional orthologs amongst complex homologous gene families. To explore the use of transcriptomic data available in public domain to identify functionally equivalent ones from all predicted orthologs, we collected genome wide expression data in mouse and rat liver from over 1500 experiments with varied treatments. We used a hyper-graph clustering method to identify clusters of orthologous genes co-expressed in both mouse and rat. We validated these clusters by analysing expression profiles in each species separately, and demonstrating a high overlap. We then focused on genes in 18 homology groups with one-to-many or many-to-many relationships between two species, to discriminate between functionally equivalent and non-equivalent orthologs. Finally, we further applied our method by collecting heart transcriptomic data (over 1400 experiments) in rat and mouse to validate the method in an independent tissue.

## 1. Introduction

Annotation of gene function is a crucial step to understand the DNA sequencing data currently generated at an unprecedented rate. The lack of functional annotation forms a major bottleneck in analyses across diverse fields, including *de novo* genome sequencing [1], Genome Wide Association Studies (GWAS) in model and non-model organisms [2], and metagenomics [3]. An experimental validation of each gene is impractical to this end as it demands high financial and time cost. It is estimated that only one percent of proteins have experimental functional annotations [4]. Bioinformatic approaches therefore provide an attractive alternative [5]. The most widely used and successful gene annotation strategy has been the annotation transfer between homologous genes. Automated annotation pipelines from sequence alone are widely used, including GOtcha [6] and BlastGO [7]. They allow fast annotation of thousands of genes for newly sequenced genomes [8]. This approach can be used within a species, where gene families (paralogs), might share common functions, or across species, where known function(s) of a gene in one species are used to infer functions of the homologous gene(s) in another species.

Despite being widely used, fast computational annotation comes at a cost of misannotation, which is present at high levels (over 10 percent) and is believed to be increasing [9] due to misannotation transfer. The most common misannotation is over-annotation, where a gene is assigned a specific but incorrect function [10]. This is partly because one of the major challenges in functional annotation transfer across species is that the orthology relationships are not always one-to-one. Specifically, a single gene in one species can be homologous to multiple paralogs in another (one-to-many homologies), after gene duplication or gene loss event(s). After a gene duplication, the two paralogs can have redundant functions, and thus should share similar functional annotations, or one copy might diverge (lose functionality, or gain new functionalities, or change cellular localisation or tissue specificity), and thus paralogs should have different functional annotations despite their homology. Similarly, multigene families (with many-to-many homologies) are highly prone to over-annotation errors.

Protein structure information can act as source for functional distinction within multigene family proteins [4]. Protein-protein interaction networks have also been successfully used to identify functional orthologs [11]; two orthologs interacting with the same proteins in each species are likely to share similar functions. Similar strategy has been applied to biochemical pathway information [12]. Co-expression gene networks have also been used in this context [13, 14, 15], as they offer two main advantages over proteinprotein interactions and biochemical pathways. First, they can be inferred from transcriptomic datasets, which are more abundant than protein-protein interaction datasets. Second, they allow functional annotation of the various classes of RNA genes. We have previously shown that multi-species information improves gene network reconstruction [16].

In order to further explore the potential of co-expressed gene networks to identify functional equivalents in complex homologous families, we collected transcriptomic data from mouse and rat liver samples. To minimise technical variation, we collected datasets generated using a single microarray platform in each species, resulting into 920 experiments in mouse and 620 experiments in rat. We firstly identified clusters of co-expressed genes using hierarchical clustering and found biologically relevant clusters. We applied an hyper-graph clustering method, SCHype [17] to simultaneously cluster co-expressed orthologous genes between species. We then focussed on 18 complex (one-to-many or many-to-many) homology groups, where at least one member in mouse and in rat where present in similar co-regulated gene clusters providing an independent source of evidence for shared functionality amongst orthologous genes in complex homologous families. We successfully applied the same method on heart transcriptomic data from mouse and rat, and investigated functional relevance of 11 other orthologous groups. Our results show the potential of this method to use co-expression as an independent measure to evaluate shared functionality amongst orthologs and limit over-zealous annotation transfers.

## 2. Methods

### 2.1. Data collection and normalisation

Microarray data for liver and heart samples in mouse and rat were collected from GEO, where data for mouse was generated using Affymetrix Mouse Genome 430 2.0 Array, and data for rat was generated using Affymetrix Rat Genome 230 2.0 Array as they were the platforms with a large number of experiments available for each species. Liver experiments came from 62 (mouse) and 28 (rat) independent studies or GEO series. Heart experiments came from 20 (mouse) and 19 (rat) independent studies or GEO series. The GEO accession numbers for individual studies are provided in supplementary table 1. Processed data was not directly comparable between studies, as different studies used different normalisation methods, leading to different distribution of values (supplementary figure 1, A and B, supplementary figure 3, A and B). As some datasets had a trimmed lower quartile for reduction in noise by limiting the variability of lowly expressed genes, we applied lower quartile trimming on all datasets (supplementary figure 1, C and D, supplementary figure 3, C and D). Specifically, we set the expression value of all probes belonging to the lower quartile to the value of the 25 percentile. We then applied quantile normalisation resulting into a uniform distribution of values for each experiment. To facilitate the comparison between mouse and rat data, we used liver mouse data as a target for quantile normalisation of heart mouse data and liver and heart rat data, using preprocessCore functions `normalize.quantiles.determine.target` and `normalize.quantiles.use.target` [18]. Liver mouse data was selected as the target because it contained more experiments than the liver rat dataset. Thus, after our normalisation steps, the distribution of values was identical for each experiment in both species.

### 2.2. Data clustering

We selected genes with variable expression across experiments by selecting probes with a standard deviation greater than one across experiments. As shown in figure 1, such probes included genes of low as well as high expression levels, and largely excluded probes showing very low expression in all experiments. Microarray data being already log-transformed, log fold change over the average values were obtained by subtracting the mean expression of each probes.

**Figure 1:**
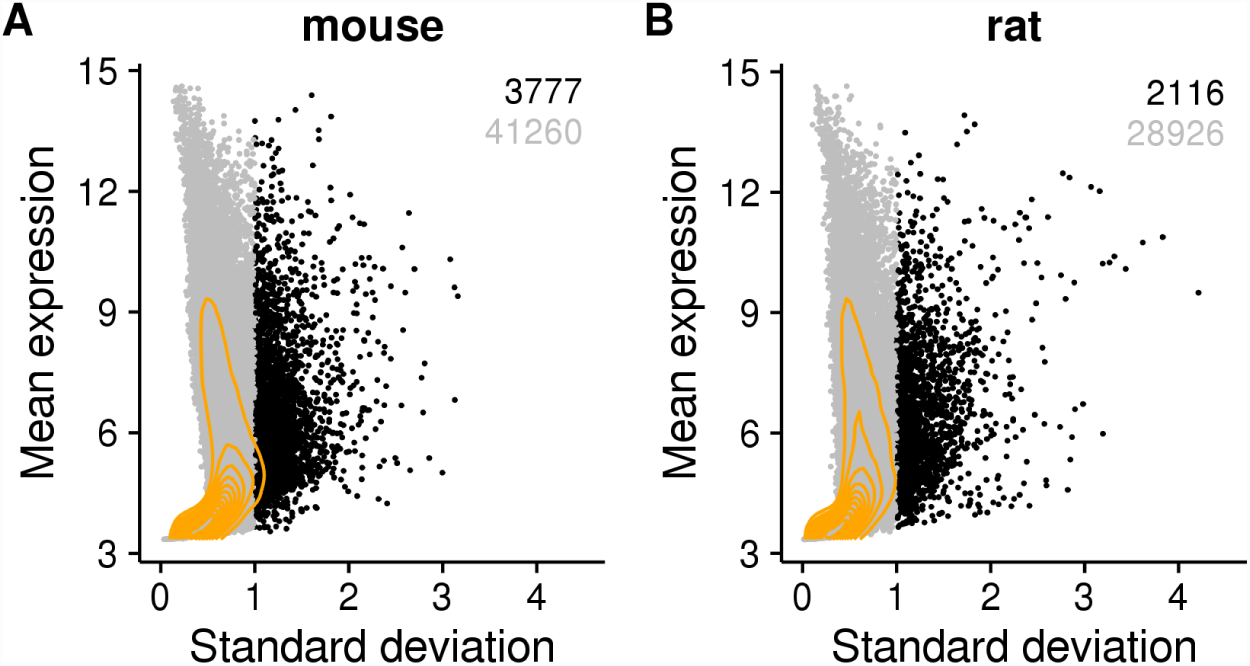
Identification of variable probes in mouse (A) and rat (B) datasets. Each dot represents a single probe. X axis: standard deviation across experiments. Y-axis: mean expression values across experiments (in arbitrary units). In black the probes with a standard deviation ≥ 1, in grey the probes with a standard deviation *<* 1. Orange lines: 2D kernel density.

Hierarchical clustering was done on the log fold change matrices using R functions `dist` ad `hclust` with default parameters (euclidean distance, complete linkage). Dendrogram branches were reordered using the function `order.optimal` from the cba package [19]. Both rows (probes) and columns (experiments) were clustered using this approach.

Gene homology information was retrieved from the Homologen database [20], and probe orthology information was obtained using the R package annotationTools [21]. Due to one-to-many homologs, rat probes and mouse probes intersections resulted into slightly different numbers for each species. Average of the two numbers was used to obtain Jaccard indexes. Jaccard index significance was obtained using the hypergeometric test, and p-values were corrected for multiple testing using Bonferroni correction.

SCHype takes as input a list of conserved interactions which was generated as follows. First Spearman correlation coefficient between each pair of probes was obtained independently for both Mouse and Rat expression data. Pairs of probes with a correlation coefficient greater or equal to 0.5 were selected. Then if orthologs of two connected probes were connected in the other species, they were kept as an SCHype input. SCHype was run using default parameters. In liver, SCHype identified 132 clusters of homologous genes co-expressed both in mouse and in rat, which included 825 nodes in mouse and 778 nodes in rat. SCHype allows probes to be included in multiple clusters. The different number of probes in mouse and rat is due to the presence of one-to-many and many-to-many orthologs, as well as the presence of gene measured by multiple probes on the array.

### 2.3. Gene ontology analysis

Gene ontology analysis was performed using PantherDB [22], using as a control gene set the genes analysed by the microarray, or only the variable gene sets previously defined.

### 2.4. Scripts and data availability

R scripts used for this analysis are available in a Github repository https://github.com/gdevailly/liver_mouse_rat. Normalised expression matrices, fold change matrices, as well as probe clusters (hierachical clustering and SCHype clustering) are available through two Zenodo collections: https://zenodo.org/record/439483 (liver data) and https://zenodo.org/record/839015 (heart data).

## 3. Results

### 3.1. Identification of variable genes across datasets

We downloaded 920 and 620 experiments for gene expression data in rat and mouse liver from the GEO database. We firstly normalised the data using lower quartile trimming (supplementary figure 1, C and D) and quantile normalization (supplementary figure 1, E and F) independently for each species. We then selected the probes with dynamic expression across samples (standard deviation ≥ 1). This resulted into 3777 probes in mouse (8.4%) and 2116 probes in rat (6.8%), with a wide range of expression values (figure 1). 735 mouse variable probes out of 3777 had a homologue in rat variable probes, and 624 rat variable probes out of 2116 had a homologue in mouse variable probes. Variable genes were enriched for pathways and functions related to liver biology (table 1), including metabolism of lipid an protein (rat, adjusted P value ≤ 10^−4^), regulation of cholesterol biosynthesis by SREBP (mouse and rat, respectively adjusted P value ≤ 0.01 and ≤ 0.03), synthesis of bile acid and salt via 24-hydroxycholesterol (rat, adjusted P value ≤ 0.03), and fatty acid metabolic process (mouse and rat, respectively adjusted P value ≤ 10^−4^ and ≤ 0.03). As the biological processes enriched in variable genes reflected functions associated with liver, we concluded that the expression variability across samples was due to biological variability, and not only technical variations, and therefore was of significance for further investigation.

**Table 1:**
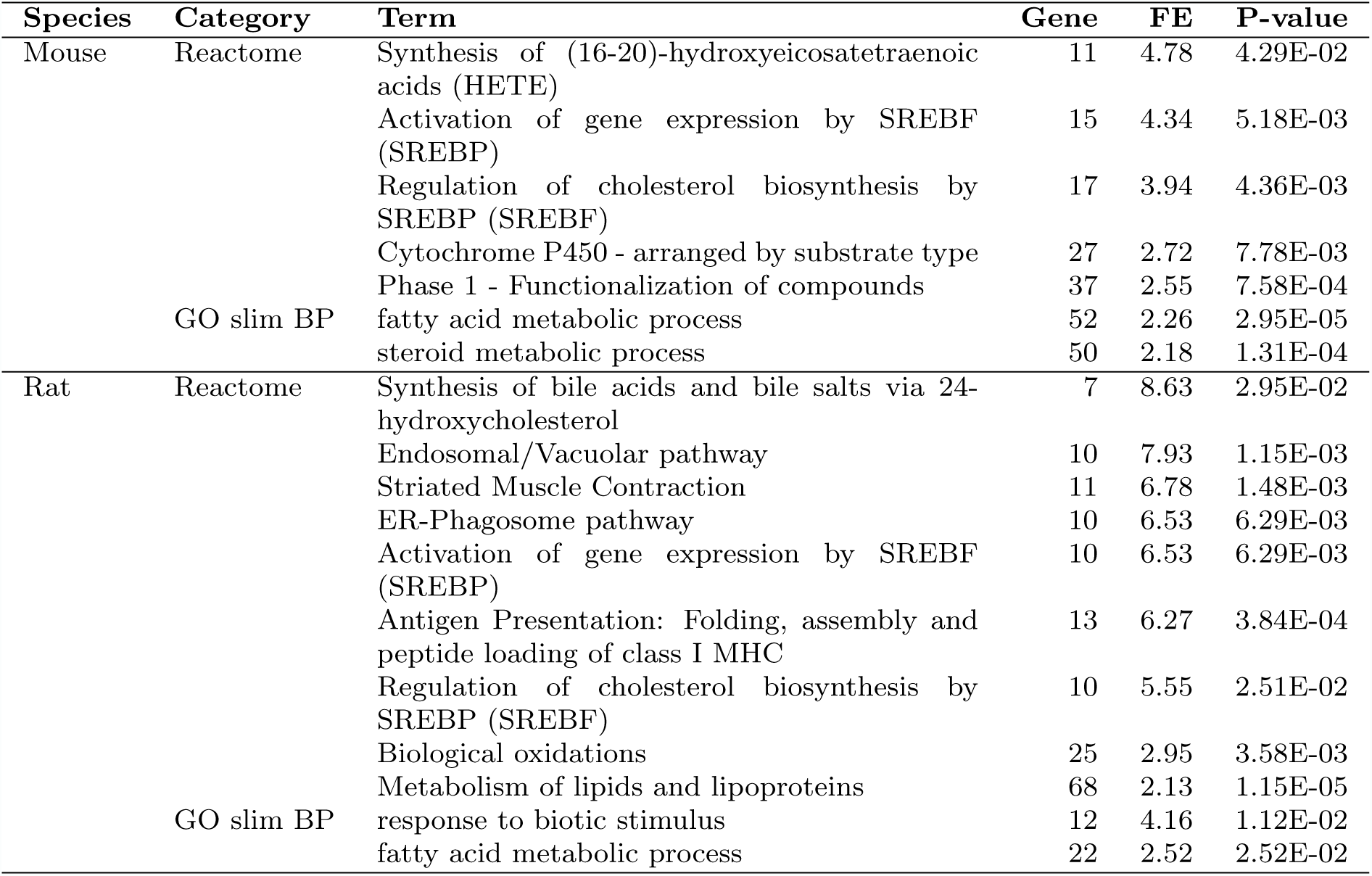
Variable genes are enriched for categories and pathways related to liver functions. FE: Fold enrichment between actual over expected number of genes. GO: gene ontology. BP: biological process. Only categories with a fold enrichment greater than 2 are shown. All P-values were corrected for multiple testing with the Bonferroni method.

### 3.2. Independent hierarchical clustering of mouse and rat data

Hierarchical clustering was applied to the mouse and rat expression matrices independently (figure 2, A and B). We defined 7 major clusters of variable probes, while the experiments were grouped in 4 clusters. The two major clusters of experiments in mouse showed broadly opposite expression patterns (figure 2A). Two major experimental groups were also noted in rat, albeit to a lesser extent compared to mouse (figure 2B). Experiments were annotated according to their series of origin (figure 2A and B, bottom of the heatmap), revealing that most experiments from the same series grouped together (including cases and controls). Notably, no series of experiments were split in the two main experiment clusters.

**Figure 2:**
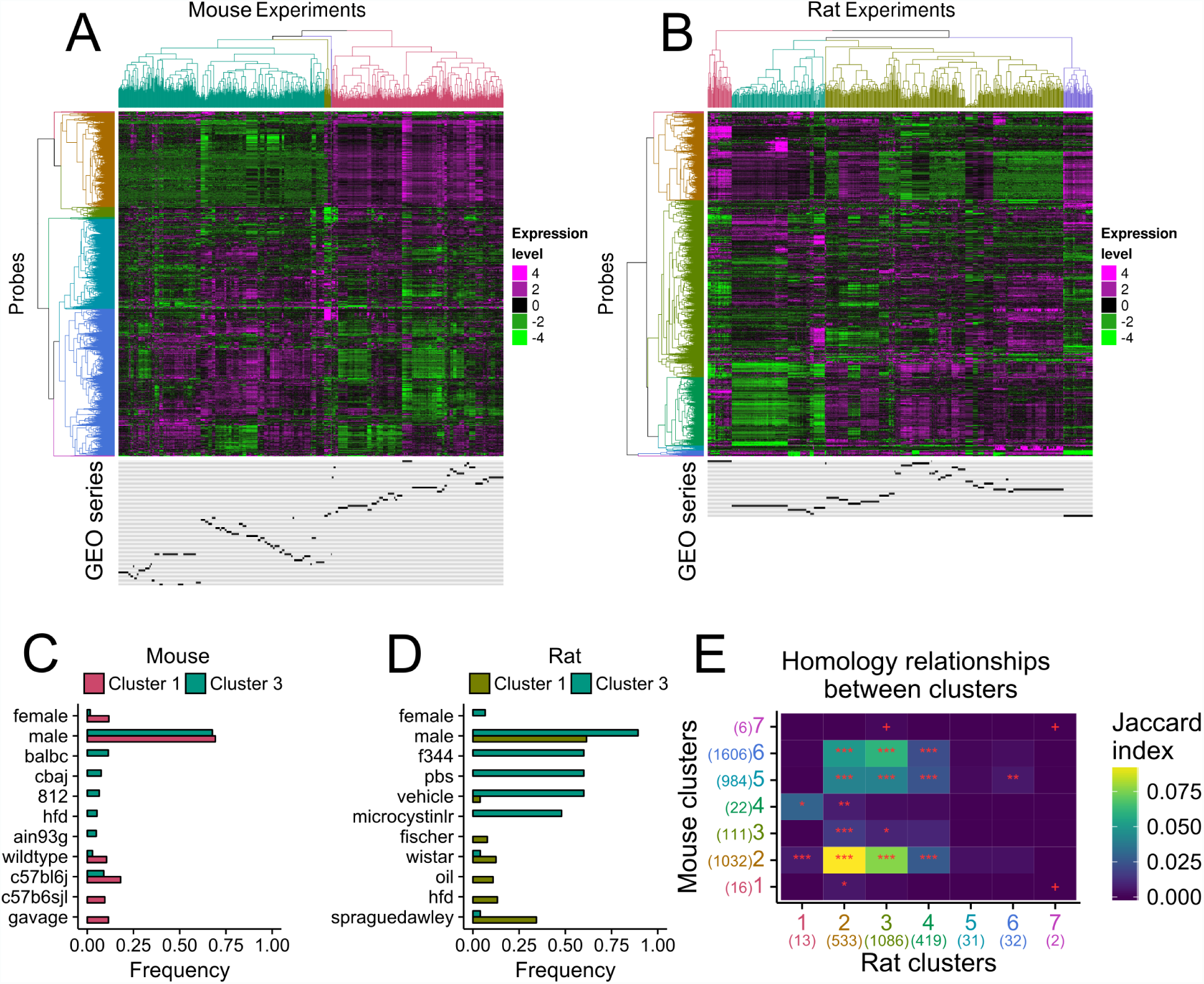
Hierarchical clustering of variable probes in mouse (A) and in rat (B). Four clusters were defined for experiments and seven for probesas reflected by the dendrogram colours. Below the heatmaps, localisation of experiments from each series are shown in black, one line per series. FC: fold change. C and D. Metadata term frequencies of the two biggest experiment clusters were compared for mouse (C) and rat (D). Colour-code matches the experiments trees in panels A and B. E. Homology relationships between probe clusters between rat (x-axis) and mouse (y-axis). Cell colour: jaccard index. Cell label: Bonferonni adjusted p-values: *** ≤ 0.0001, ** ≤ 0.001, * ≤ 0.01, + ≤ 0.05. Cluster number colours match the probes dendrogram colours in panels A and B. The numbers in parenthesis denote the number of probes in each cluster.

We characterised the main experiment clusters by looking at the most different non-trivial terms in the element-term matrix build from the meta-data retrieved from GEO (characteristic field, figure 2, C and D). No clear difference between experiment clusters was observed in mouse. Experiment cluster 3 in rat seems to be composed mostly of F344 strains of rat and/or of rat treated with the microcystinlr toxin. To note, this cluster is dominated by experiments from a single experiment series (figure 2B). Since experiment clustering matched series of origin of the data, this hinders correction for batch effects to get biological differences.

Given that mouse and rat probes formed two major clusters anti-correlated with each other despite diverse experimental set ups in each species, we investigated whether the mouse and rat probe clusters were composed of probes measuring similar genes (figure 2E). We calculated the overlap between genes in each cluster in mouse with genes in each cluster in rat. Cluster 2 in mouse (golden colour, figure 2A) and cluster 2 in rat (golden colour, figure 2B) showed a very high overlap with the highest Jaccard index across all clusters. Neither mouse cluster 2 nor rat cluster 2 were enriched for any gene ontology term or reactome pathway terms, when using the set of variable probes as background. Most clusters did not show a very high genes overlap across species. This might be due to the fact that the experiments carried out in each species were different, resulting in distinct set of genes perturbed in each species, resulting into little overlap of co-expression clusters across species. Functional enrichment analyses of other clusters were suggestive that observed gene variations reflected differences in the liver physiology. Specifically, cluster 1 in mouse (claret red colour, figure 2A) was enriched for generation of precursor metabolites and energy (adjusted P value ≤ 10^−6^), steroid metabolic process (adjusted P value ≤ 0.001), fatty acid metabolic process (adjusted P value ≤ 0.001), and Cytochrome P450 arranged by substrate type (adjusted P value ≤ 10^−6^). Cluster 3 in mouse (green colour) was enriched for arachidonic acid metabolic process (adjusted P value ≤ 0.01), icosanoid metabolic process (adjusted P value ≤ 0.05), fatty acid derivative metabolic process (adjusted P value ≤ 0.05), and Cytochrome P450 arranged by substrate type (adjusted P value ≤ 0.05). Cluster 6 in rat (blue) was enriched for proteolysis (FE 10, adjusted P value ≤ 0.01). More terms related to the liver matabolism were enriched when the same analysis was performed using all genes as a background (supplementary table 2).

### 3.3. Co-clustering of Mouse and Rat expression data

To identify clusters of homologous probes between mouse and rat, we used the hyper-graph clustering tool SCHype [17]. SCHype uses a recur-sive spectral clustering algorithm to identify sets of nodes in each species with a greater than expected number of conserved interactions (based on co-expression in this case) between them (figure 3A). Input data for SCHype was built using three graphs: a mouse probe graph built from pairs of probes with a Spearman correlation coefficient ≥ 0.5 (supplementary figure 2A), a rat probe graph with pairs of probes with a Spearman correlation coefficient ≥ 0.5 (supplementary figure 2B), and a probe to probe homology graph between rat and mouse built using the Homologene database [20] and the annotationTools package [21]. SCHype identified 132 clusters of homologous genes co-expressed both in mouse and in rat, which included 825 nodes in mouse and 778 nodes in rat (figure 3B). SCHype allows probes to be included in multiple clusters resulting into 474 unique probes in mouse and 425 unique probes in rat. It identified four clusters with over 30 homologous genes in each species, eighteen clusters with over 10 probes in each species, thirtyfive clusters with only 2 co-expressed probes in each species (figure 3B). We further focussed on the first four (c1-c4) SCHype clusters (figure 3C). We firstly compared SCHype clusters with results obtained by clustering data from each species independently. SCHype cluster c3 highly overlapped with the previous cluster 2 in mouse (golden colour, figure 2A) and the cluster 2 in rat (golden colour, figure 2B). These two clusters were shown to share a high number of homologous probes (figure 2E). Gene ontology analysis of the four biggest SCHype clusters, both over the set of variable probes or over the full set of probes, did not lead to any significant results, most likely due to small number of genes in each cluster. Importantly, the experiments in each series no longer clustered together after restricting the data to each of the four biggest SCHype clusters (figure 3C). Individual experiments from each series nevertheless belonged to the same large experiment cluster (figure 3C) highlighting the need for building an expression compendium to obtain these results.

**Figure 3:**
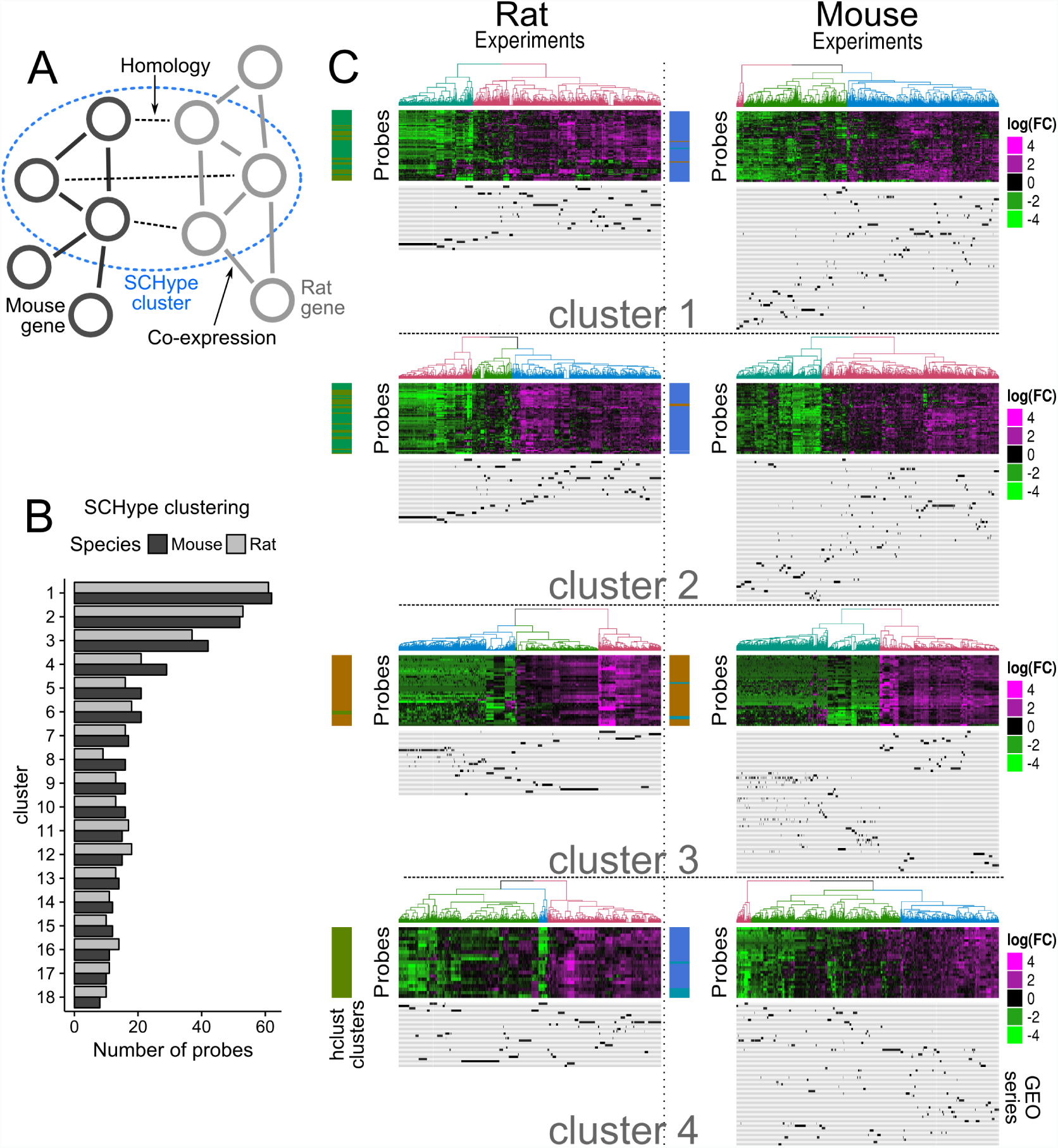
Co-clustering of rat (middle) and mouse (right) liver data using SCHype. A. SCHype is a clustering tool for hyppergraphs, built here from two co-expression graphs and an homology graph. B. Number of mouse (dark grey) and rat (light grey) probes for the SCHype clusters with more than 10 probes for each species. X-axis: Number of probes included in each SCHype cluster. Y-axis: SCHype predicted clusters, numbered according to the number of probes per cluster in decreasing order. C. The biggest four SCHype clusters are shown. Genes in mouse and rat in each cluster are homologous to each other. The results of hierarchical clustering for each species is shown as a colour bar on the left. colour-code matches the experiments trees in figure 1. Under the heatmap, clustering localisation of experiments from each series are shown in black, one line per series.

### 3.4. Co-clustering across species as source of information for inferring shared functionality amongst orthologs

SCHype clustering successfully identified clusters of homologous genes co-expressed in both mouse and rat datasets. This information adds an independent evidence in support of a functional annotation transfer for pairs of orthologous genes across species found in the same SCHype cluster(s), as functionally equivalent orthologs would be co-expressed with the same set of genes in both species, and therefore would be included in the same SCHype cluster(s). We investigated if SCHype clusters could help identify functionally equivalent orthologs amongst complex homology groups. Eighteen homology groups of three members or more had at least one member of each species in the same SCHype cluster(s). For example, for homology group 137299 (table 2), *Anp32a* in mouse and *Anp32a* in rat were in the same SCHype cluster 69, while *LOC100909983*, another homologue of rat *Anp32a*, was not. This suggests that indeed *Anp32a* in rat is the functional equivalent of *Anp32a* in mouse, but *LOC100909983* is not. In this case, our method found back a functional equivalent already known [23]. Similar observations were made for homology groups 68982 (*Ccnb1*), 10699 (*Cdc248*), 3938 (*Ppp1r3c*), and 14108 (*Rasl10b*) (table 2). In five cases, all members of the homology groups were included in the same SCHype clusters (table 3), suggesting that all orthologs are likely to share the same function(s). Finally, eight homology groups showed more complex situations, where neither only one nor all the homologs where present in the same groups (table 4 and supplementary table 3). For example, in homology group 117945, *Cyp2c7* in rat had three homolous genes in mouse but only *Cyp2c38* in mouse belonged to the same SCHype cluster (table 4) predicting that mouse *Cyp2c38* (and not mouse *Cyp2c29* or mouse *Cyp2c39*) is a functional ortholog of rat *Cyp2c7*. We further explored the impact of the correlation threshold used to build the hypergraph (0.5, supplementary figure 2) on the functional transfer evidence generated by assessing the predictions made using a higher correlation threshold of 0.75 (supplementary table 3). As expected, this resulted in reduction of co-expression edges, and thus reduction in identified clusters. Of the 18 groups described, 6 retained at the threshold of 0.75, with no major changes on the predictions of shared functionality. Altogether, hypergraph clustering of co-expression network from rat and mouse liver microarray data was able to provide new evidence for functional annotation transfer between orthologous groups.

**Table 2:** SCHype clustering of homologous groups: SCHype gene clustering reflects gene names. Homology groups were obtained from the Homologene database. Tick mark indicates the inclusion of the gene in the corresponding SCHype cluster.

| Homology group | Species | Gene name | SCHype cluster |  |
| --- | --- | --- | --- | --- |
| 137229 |  |  | cluster 69 |  |
|  | mouse | *Anp32a* | ✓ |  |
|  | rat | *Anp32a* | ✓ |  |
|  | rat | *LOC100909983* |  |  |
| 68982 |  |  | cluster 7 | cluster 30 |
|  | mouse | *Ccnb1* | ✓ | ✓ |
|  | mouse | *Gm5593* |  |  |
|  | rat | *Ccnb1* | ✓ | ✓ |
| 10699 |  |  | cluster 2 | cluster 118 |
|  | mouse | *Cd248* | ✓ | ✓ |
|  | rat | *Cd248* | ✓ | ✓ |
|  | rat | *LOC100911932* |  |  |
|  | rat | *LOC100911882* |  |  |
| 3938 |  |  | cluster 1 |  |
|  | mouse | *Ppp1r3c* | ✓ |  |
|  | rat | *Ppp1r3c* | ✓ |  |
|  | rat | *LOC100910671* |  |  |
| 14108 |  |  | cluster 2 |  |
|  | mouse | *Rasl10b* | ✓ |  |
|  | rat | *Rasl10b* | ✓ |  |
|  | rat | *LOC100912246* |  |  |

**Table 3:** SCHype clustering of homologous groups: all members of the homology groups share predicted functionalities. Homology groups are obtained from the Homologene database. Tick mark indicates the inclusion of the gene in the corresponding SCHype cluster.

| Homology group | Species | Gene name | SCHype cluster |  |  |
| --- | --- | --- | --- | --- | --- |
| 128630 |  |  | cluster 9 | cluster 12 | cluster 45 |
|  | mouse | *Ceacam1* | ✓ |  |  |
|  | mouse | *Ceacam2* | ✓ | ✓ | ✓ |
|  | rat | *Ceacam1* | ✓ | ✓ | ✓ |
| 11456 |  |  | cluster 5 |  |  |
|  | mouse | *Elovl6* | ✓ |  |  |
|  | rat | *Elovl6* | ✓ |  |  |
|  | rat | *LOC102549542* | ✓ |  |  |
| 20277 |  |  | cluster 35 |  |  |
|  | mouse | *Rrm2* | ✓ |  |  |
|  | rat | *Rrm2* | ✓ |  |  |
|  | rat | *LOC100359539* | ✓ |  |  |
| 55991 |  |  | cluster 1 | cluster 119 |  |
|  | mouse | *Tmed2* | ✓ | ✓ |  |
|  | mouse | *Gm21540* | ✓ | ✓ |  |
|  | rat | *Tmed2* | ✓ | ✓ |  |
| 11890 |  |  | cluster 10 | cluster 43 | cluster 81 |
|  | mouse | *Tnks2* | ✓ | ✓ | ✓ |
|  | rat | *LOC100910717* | ✓ | ✓ | ✓ |
|  | rat | *Tnks2* | ✓ | ✓ | ✓ |

**Table 4:** SCHype clustering of homologous groups: new predictions for functional orthologous relations. Homology groups are obtained from the Homologene database. Tick mark indicates the inclusion of the gene in the corresponding SCHype cluster. Four additional, more complex, homology groups are shown in supplementary table 3.

| Homology group | Species | Gene name | SCHype cluster |
| --- | --- | --- | --- |
| 117948 |  |  | cluster 102 |
|  | mouse | *Cyp2c38* | ✓ |
|  | mouse | *Cyp2c29* |  |
|  | mouse | *Cyp2c39* |  |
|  | rat | *Cyp2c7* | ✓ |
| 104115 |  |  | cluster 33 |
|  | mouse | *Hsd3b5* | ✓ |
|  | mouse | *Gm10681* |  |
|  | mouse | *Hsd3b4* |  |
|  | mouse | *Gm4450* |  |
|  | rat | *Hsd3b5* | ✓ |
|  | rat | *LOC100911116* | ✓ |
| 137425 |  |  | cluster 2 |
|  | mouse | *Lce3c* | ✓ |
|  | rat | *LOC100361951* | ✓ |
|  | rat | *LOC100911982* | ✓ |
|  | rat | *Lce3d* |  |
| 129514 |  |  | cluster 17 |
|  | mouse | *Rdh9* | ✓ |
|  | mouse | *Rdh1* |  |
|  | mouse | *Rdh16* |  |
|  | mouse | *Rdh19* |  |
|  | mouse | *BC089597* |  |
|  | rat | *Rdh16* | ✓ |
|  | rat | *LOC100365958* | ✓ |

### 3.5. Functional annotation transfer across rat and mouse using heart transcriptomic data

To test whether the approach described above was extendible to other tissues, we collected expression data in heart for mouse and rat. Data from heart samples processed using the same microarray platform were downloaded from GEO, for a total of 248 experiments from 20 studies in mouse and 1202 experiments from 19 studies in rat. Same data processing pipeline as for the liver samples (supplementary figure 3) resulted into selection of 7371 (mouse) and 917 (rat) variable genes for clustering (supplementary figure 4). The large difference between the selected number of genes might be due to the large difference in the number of samples available (and therefore used in analysis) for each species. Functional enrichment analysis of the variable genes revealed pathways related to the heart functions (supplementary table 4), confirming that at least part of the gene expression variability was reflecting biological differences. Hierarchical clustering of the mouse and rat dataset separately revealed clusters of probes and clusters of experiments (supplementary figure 5). Notably, experiments from a same series (GEO) were split in distinct clusters.

**Figure 4:**
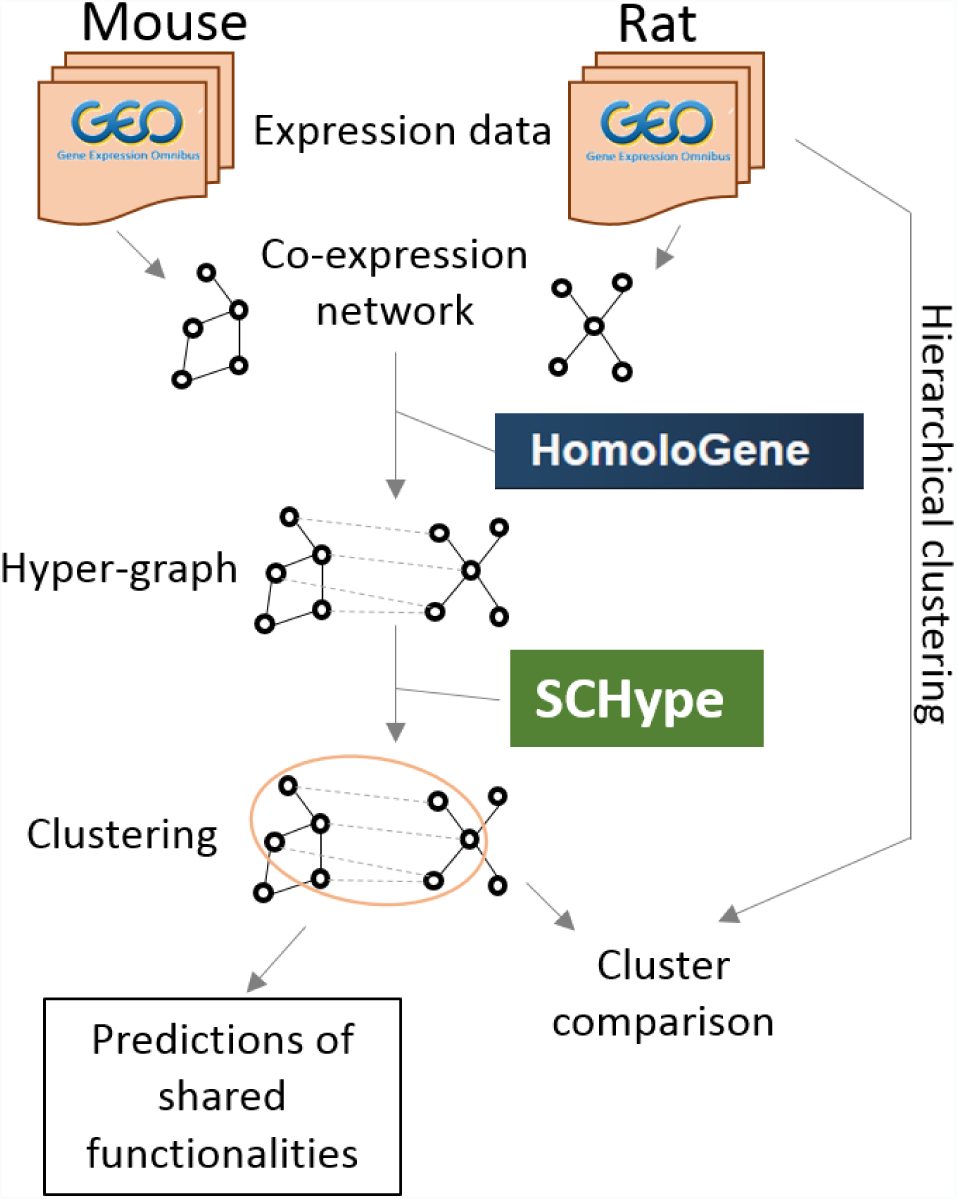
Work-flow diagram. Transcriptomic data (microarray) was gathered from GEO to build species-specific co-expression network. Homology information from the Homologene database together with co-expression networks were used to extract hypergraph clusters using the SCHype software. Resulting clusters were firstly compared to species specific hierarchical clusters, and were used to infer shared functionality links in complex homology groups.

Co-clustering of the mouse and rat co-expression network in heart, using Homologene homology information and the SCHype hypergraph clustering tool, identified clusters of homologous genes co-expressed in both species (supplementary figure 6). Notably, this provided an independent evidence in favour or against shared functionality for 12 complex orthology groups (supplementary table 5). In details, for 6 groups all homologous genes where found in the same SCHype cluster (*i.e.* were in homologous co-expressed gene networks in both species). For 3 one-to-two homology groups (*Ifit3*, *Ogn* and *Ppp1r3c*), only one of the two paralogs was included in the same SCHype cluster(s) as the ortholog copy, suggesting a loss or change of functionality for the second paralog. The remaining 3 groups presented complex scenarios where more than one, but not all paralogs were included in the same SCHype cluster(s). One of these complex groups, Homologene group 128352, was found both in heart and liver data. Altogether, the proposed method was able to provide evidence to support annotation transfer from transcriptomic data not only in liver, but also in heart, suggesting that the approach is applicable to different tissues.

## 4. Discussion

Here we have shown that transcriptomic data can be used to provide evidence for functional annotation transfer between othologs, using co-expression networks built from mouse and rat liver and heart samples. In liver, we identified 18 complex homologous groups (i.e. with paralogs in at least one of the species), including 54 genes in mouse and 46 genes in rat, with at least one gene in mouse and one gene in rat in the same SCHype cluster(s). Twelve more groups (of which 11 groups non overlapping with the liver analysis) were found when applying the same method to heart transcriptomic data. Increasing the correlation thresholds resulted in loss of total number of predictions, as expected. Lowering the correlation threshold and the standard deviation threshold, on the other hand, will likely increase the number of homologous groups, potentially with a higher false positive rate. The use of co-expression network to provide evidence for functional annotation transfer has been previously demonstrated [13, 14, 15]. These studies combined samples from various tissues, while we analysed two tissues (liver and heart) independently. Both approaches are complementary. While mixing tissues might result in broader co-expression network (many edges), it might also lack the fine resolution needed to improve functional annotation inference in a tissue specific context. We used microarray data in this study as it is by far the most abundant dataset. However, consortia like GTEx [24] have generated large amount of RNA sequencing data, and we envisage application of the method described here to RNA sequencing data in the future. The greater sensitivity of RNA sequencing over microarray [25] might allow the identification of more co-expressed genes.

Despite rigorous data normalisation, liver experiments from the same series tended to cluster together, cases and controls included. While this could be a sign of technical biases, gene ontology analysis of the variable genes demonstrated that they are related to liver functions. Thus it appears that the gene expression variability we observed is, at least partially, reflecting biological variations impacting the liver physiology. Importantly, individual experiments from series did not cluster together in SCHype clusters. We applied various approaches but could not identify the biological origin(s) of the observed variations. This is in part due to the lack of standardised experiment metadata fields in GEO (not all datasets even had a strain or a sex information, for example), and the lack of controlled vocabulary used to describe experiments. It is a possibility that better annotation of the metadata would have allowed the identification of critical confounding factors. Noteworthy, heart experiments were not clustered by series of experiments. It could be that the heart tissue is less sensitive than the liver to differences in the animal environment.

SCHype clustering was able to find some known as well as some yet to be experimentally validated ortholog functional relationships. For example, only mouse and rat *Ccnb1* were in the same SCHype cluster, and not *Gm5593*. While mouse *Ccnb1* and rat *Ccnb1* are annotated as protein coding genes, *Gm5593* in mouse is annotated as a processed pseudogene [23].

Finally, we note that conserved co-expression of orthologous genes is not a direct proof of shared functionality, but it can be used as an additional layer of evidence. While protein-protein interaction networks could be used for the same aim, transcriptomic data are more easily generated and therefore more likely to be widely available for many species. Thus the method described here shows a promise to enhance functional gene annotation transfer across species. It can provide an experimental support for one-to-one ortholog annotation transfer, and can help identify functionally similar and non similar orthologs in one-to-many and many-to-many orthology groups.

